# Hypoxia classifier for transcriptome datasets

**DOI:** 10.1101/2021.11.15.468572

**Authors:** Laura Puente-Santamaría, Lucia Sanchez-Gonzalez, Ricardo Ramos-Ruiz, Luis del Peso

## Abstract

Molecular gene signatures are useful tools to characterize the physiological state of cell populations, but most have developed under a narrow range of conditions and cell types and are often restricted to a set of gene identities.

Focusing on the transcriptional response to hypoxia, we aimed to generate widely applicable classifiers sourced from the results of a meta-analysis of 69 differential expression datasets which included 425 individual RNA-seq experiments from 33 different human cell types exposed to different degrees of hypoxia (0.1-5%O_2_) for 2-48h.

The resulting decision trees include both gene identities and quantitative boundaries, allowing for easy classification of individual samples without control or normoxic reference. Each tree is composed by 3-5 genes mostly drawn from a small set of just 8 genes (EGLN1, MIR210HG, NDRG1, ANKRD37, TCAF2, PFKFB3, BHLHE40, and MAFF). In spite of their simplicity, these classifiers achieve over 95% accuracy in cross validation and over 80% accuracy when applied to additional challenging datasets. Our results indicate that the classifiers are able to identify hypoxic tumor samples from bulk RNAseq and hypoxic regions within tumor from spatially resolved transcriptomics datasets. Moreover, application of the classifiers to histological sections from normal tissues suggest the presence of a hypoxic gene expression pattern in the kidney cortex not observed in other normoxic organs. Finally, tree classifiers described herein outperform traditional hypoxic gene signatures when compared against a wide range of datasets. This work describes a set of hypoxic gene signatures, structured as simple decision tress, that identify hypoxic samples and regions with high accuracy and can be applied to a broad variety of gene expression datasets and formats.

## Introduction

A gene expression signature is a single or combined group of genes whose expression is altered in predictable way in response to a specific signal or cellular status. Gene signatures are often derived from the set of differentially expressed genes (DEGs) identified when comparing two groups of transcriptomes, such as disease versus healthy controls or treated versus untreated samples. In turn, a gene signature can be of aid in trying to determine whether a given biological sample was exposed to that particular stimulus or belongs to the status defined by the gene set. Thus, reliable gene signatures can be used as surrogate markers for the activation of pathways or cellular status.

Hypoxia can be defined as the situation were oxygen supply does not meet cellular demand [1]. In response to hypoxia cells activate a gene expression program, under the control of the Hypoxia Inducible Factors (HIFs) [2], that aims to increase oxygen supply while reducing its consumption. Thus, this transcriptional response restores oxygen balance and, as such, it is central in maintaining tissue homeostasis. Importantly, oxygen homeostasis is disrupted in a number of prevalent pathologies including neoplasms [3] and cardio-respiratory diseases [4]. For all this reasons, the development of a hypoxic gene signature could be of practical interest to identify cells or samples that had been exposed to hypoxia, and accordingly, a number of studies have published hypoxic gene signatures [2, 5–11]. However, in spite of their merit, in all these cases the gene signature was derived from a limited set of related tumoral samples, raising the question of their applicability in other contexts. On another note, in almost all the cases, the gene signature is just a set of genes without any additional information reflecting their relative importance or their expected expression levels under normoxic/hypoxic conditions, meaning that it is nearly impossible to classify an individual isolated sample as normoxic or hypoxic based solely in the identities of the genes in the signature.

Herein we describe tree-based classifiers that accurately identify hypoxic cells or samples based on their gene expression profile. The identification is absolute, meaning that it does not require a set of normoxic reference samples to sort out the hypoxic ones. Thus, it can be applied to interrogate a single isolated sample. Finally, although the classifier implicitly contains information about the relative importance of the genes in the signature and their expression levels in hypoxia, it is simple enough to be interpreted and applied without the need for sophisticated computational tools.

## Materials and Methods

### RNA-seq data download and processing

Raw reads of the RNA-seq experiments were downloaded from Sequence Read Archive [12]. Pseudocounts for each gene were obtained with salmon [13] using RefSeq [14] mRNA sequences for human genome assembly GRCh38/hg38 and mouse genome assembly mm10 as references.

Reads counts of tumoral and healthy samples were downloaded from the TCGA data portal and transformed to counts per million.

Spatial gene expression datasets were downloaded from 10X Genomics website [15–21]. Raw read counts are normalized with sctransform[22] following Seurat v4.0.4[23] standard pipeline for analysis, visualization, and integration of spatial datasets.

### Generation of a classifier

To generate the classifier we made use of 425 transcriptomic profiles of hypoxiaexposed cells and their normoxic counterparts described in a recent study [24]. From the gene pseudocounts in each sample, we calculated each gene’s ranking percentile and used this information in downstream analyses. We used the R package randomForest [25] to perform feature selection and the R package rpart [26] to generate decision trees. Each decision tree was evaluated by cross-validation using 70% of the available RNA-seq experiments as a training set and the remaining 30% as a validation set. By default, a sample is classified as hypoxic when the tree assigns it a probability over 50% of being hypoxic, even though this threshold can be made stricter or laxer.

## Results

### Generation of a hypoxic classifier

Results from our previous work [24] on differential expression triggered by hypoxia indicate that, even for those genes showing a significant regulation in the ensemble of datasets, the response to hypoxia could be in large part cell-specific. Thus, we sought to identify a minimal set of genes that could be used as a reliable readout of exposure to hypoxia and to develop a simple, easy to use, classifier that could identify whether an individual sample is hypoxic based on its gene expression.

With the goal of making the model as widely applicable as possible we chose to use as input data the percentile of each gene on a gene expression ranking, thus minimizing the effects of read depth, different normalization methods, possible rRNA contamination, and other factors that influence RNA quantification. Thus, we first constructed a gene ranking matrix for a set of 425 individual RNA-seq samples derived from published transcriptomic analysis of hypoxic cells and controls (Fig. 1A). For subsequent analyses we kept the subset of 178 genes both significantly upregulated by hypoxia (*LFC* > 0.7, *FDR* < 0.01) and widely expressed (detectable in ≥ 90% of the analyzed subsets) according to a meta-analysis performed on this data set [24].

**Figure 1.**
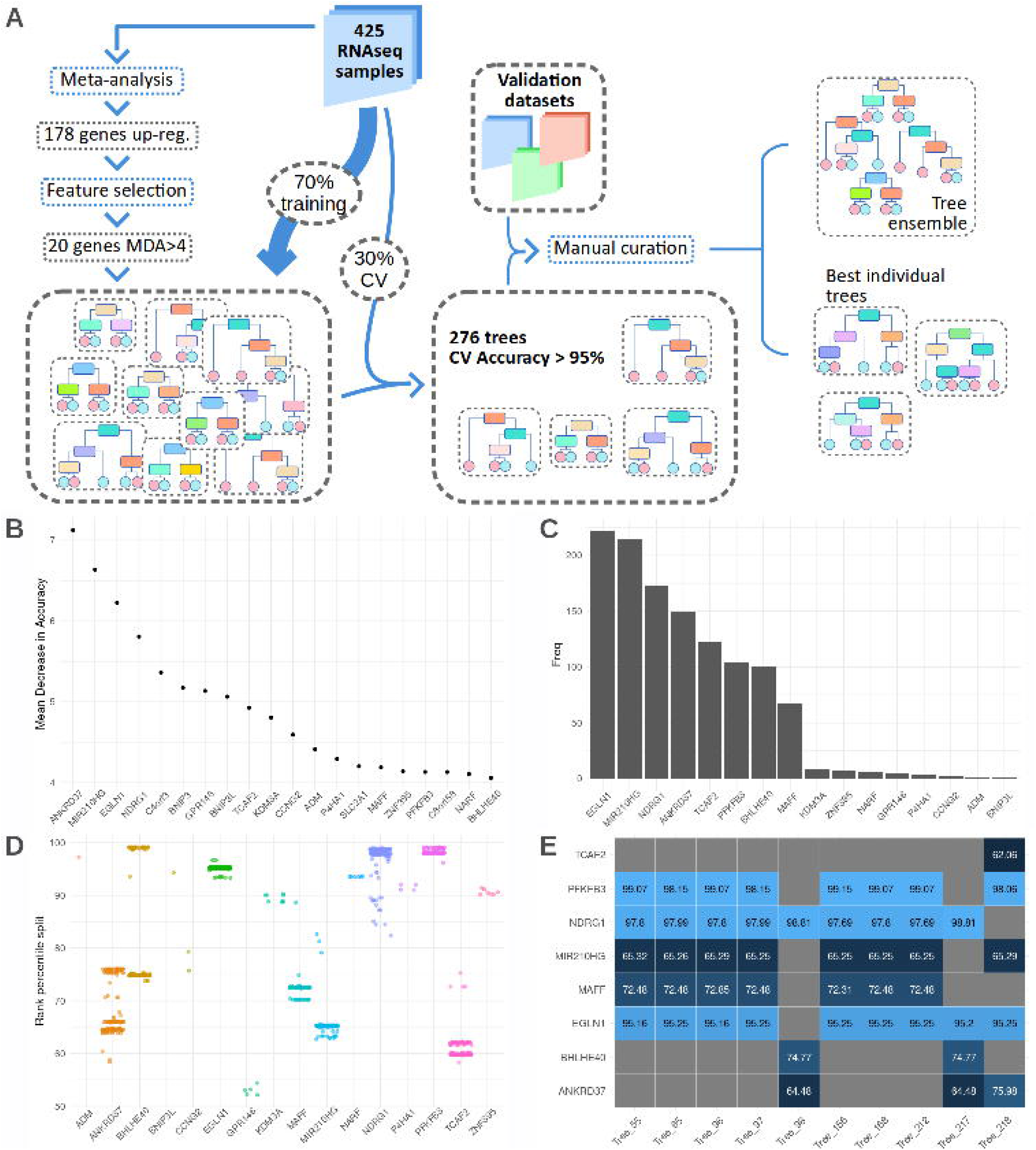
Generating expression based tree classifiers to identify hypoxic samples. **A**:Decision trees generation overview. 425 RNA-seq samples exposed to normoxia or hypoxia were processed to produce a ranking set of genes from each of one them. A subset of the resulting data matrix, consisting in the 178 genes significantly up-regulated by hypoxia according to ref [24], was used as input to a feature selection algorithm. The 20 genes showing an MDA>4 were then selected to generate 10000 random trees and the 276 trees showing an accuracy over 95% in cross validation were selected as classifiers. Finally, a set of challenging datasets not used in the generation nor training steps, were used to test the performance of the 276 trees and select the best overall tree and two additional substitutes.**B**: median decrease in accuracy index of the 20 most important genes according to 1000 random forest iterations. **C**: Frequency of each gene being used as a predictor variable in the classification trees. **D**: Split points for the rank percentile (100 being the most expressed gene, 0, the least) of the genes used in all the models with accuracy > 0.95. **E**: Split points for the rank percentile of the genes used in the 10 best performing models according to cross-validation accuracy.

In order to select the most informative genes in this subset, we used 1000 iterations of a random forest classifier sampling 70% of the RNA-seqs at each iteration (Fig. 1A). As a measure of each gene’s importance we use the mean decrease in accuracy (MDA), representing how much accuracy the model losses by excluding each gene, across all iterations. The 20 genes with average MDA over 4 (Fig. 1B) were selected to train 10000 decision trees randomly sampling 70% of the individual RNA-seq experiments and using the remaining 30% as a validation set. The 276 trees with an accuracy over 0.95 on the validation set were selected to further test their performance (Supplementary table S1-3, ”Cross validation” and Fig. 1A)).

Only 16 genes are used in all of the 276 decision trees, with half of them (EGLN1, MIR210HG, NDRG1, ANKRD37, TCAF2, PFKFB3, BHLHE40, and MAFF) being included overwhelmingly more frequently (Fig. 1C). Most of these genes have already been linked to the transcriptional response to hypoxia [27–34], even though in some cases their particular role in it has not yet been defined.

In these classification trees the rank percentile of the expression of gene the in each decision point is used to determine the branch followed for the classification of the sample, hence final sample label is assigned based on the relative (percentile rank) expression values of the genes in the tree. As seen in Fig. 1D, the split point for most genes is limited to a narrow range of rank percentiles, with ANKRD37 and BHLHE40 being the exception. These two genes show two differentiated split points that depend on the identity of remaining genes in the tree: for ANKRD37, it depends on whether its combined together with NDRG1 or TCAF2, while BHLHE40’s depends on whether the tree includes MIR210HG. Fig. 1E represents gene identity and split points for the 10 best trees according to cross-validation accuracy. Both in this top 10, as well as in the whole set of 276 trees we find a limited number of topologies present, with MIR210HG, EGLN1, MAFF, NDRG1, and PFKFB3 forming the most common combination, closely followed by ANKRD37, NDRG1,and BHLHE40.

### Evaluation and validation of the resulting decision trees

In order to evaluate the performance of each one of the 276 decision trees with an accuracy over 95% and test particular strengths and weaknesses of each model, we tested them on a series of datasets that were not part of the training nor crossvalidation sets and had some differential feature that posed a challenge to the classification (Supplementary table S1-1 ”RNA-seq metadata”). In addition to evaluate their performance, the result of these analyses guided us in the selection of the trees best suited to be used as general and robust hypoxia classifiers.

First we chose a time series experiments available on PRJNA561635 [35], consisting of a set of transcriptional profiles of Human Umbilical Vein Endothelial Cells (HUVEC) exposed to different oxygen concentrations at nine time points. The main challenge with this validation set is detecting early stages of hypoxia (1-3 hours), where most hypoxia-target genes have just barely began to accumulate, and differentiate mild hypoxic stress (3% oxygen) from physoxia (5% oxygen), which is within the range of physiological oxygen concentration found in many tissues [36] and hence *in vitro* trigger a weaker transcriptional response for many genes [37]. As shown in Fig. 2A and Supplementary table S1-4 ”PRJNA561635”, all decision trees correctly classified normoxic samples and samples exposed to oxygen levels at or below 3%O_2_ (i.e. physiological hypoxia,[36]) for at least 5h. It is worth highlighting that around a third of the threes were also able to detect earlier stages of hypoxia (2h 1%O_2_, 3h 3%O_2_). In addition, these results clearly show that the lower the oxygen tension, the strongest the signal detected at early times of exposure with 5% oxygen being at the boundary between normoxia and physiological hypoxia.

**Figure 2.**
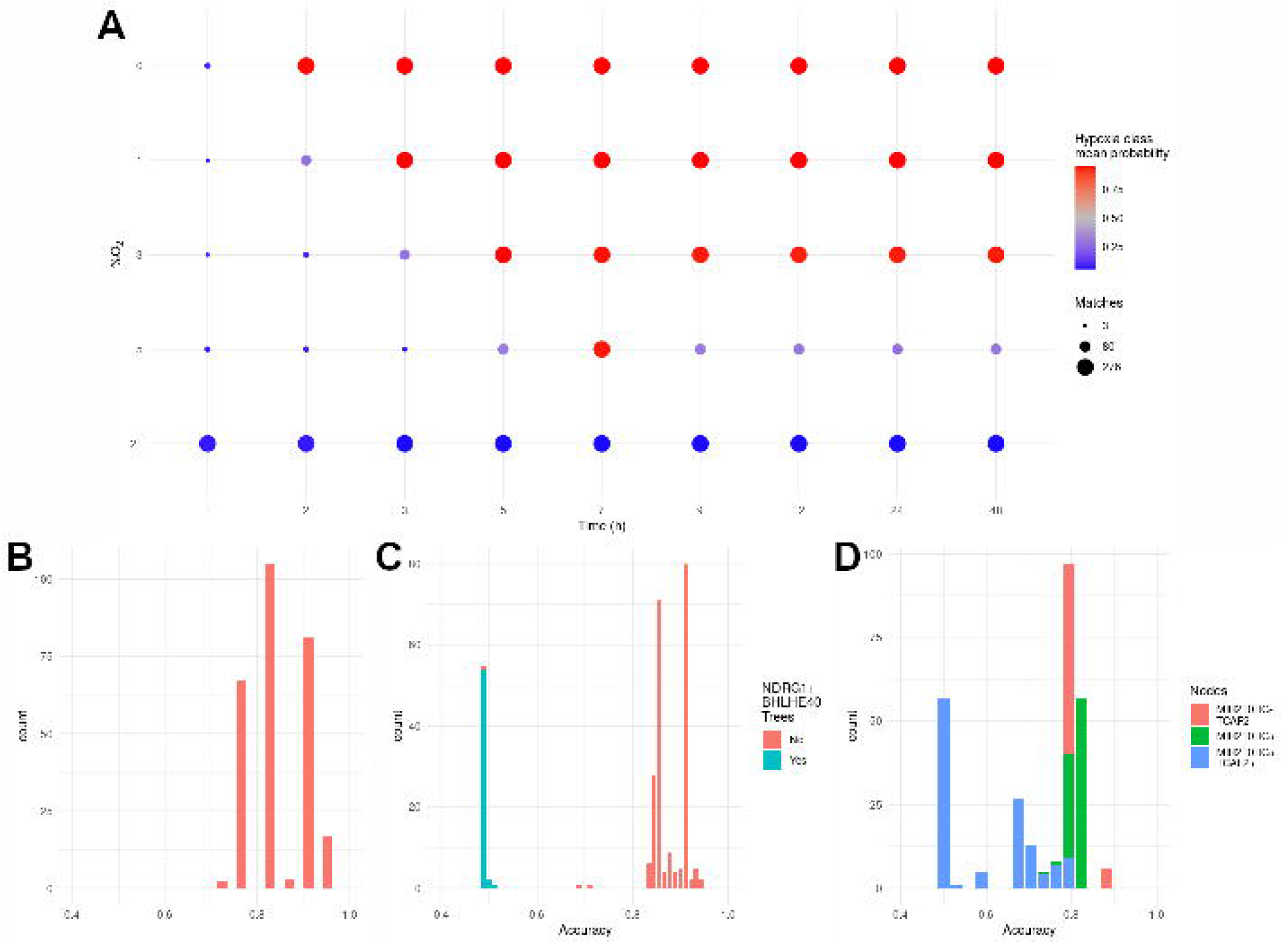
Validation of tree classifiers on novel datasets. **A:** Time and oxygen series dataset. Each of the dots represents one of the samples in the five time series in PRJNA561635, ordered by oxygen tension and time. The color of each dot represents the mean probability of each sample to be classified as hypoxic, while the size of the dot is proportional to the number of trees correctly classifying each sample. **B-D:** Distribution of the accuracy of the 276 classification trees on validation datasets. Different colors are used to identify trees including specific genes and/or topology as indicated in the legend. See text for details. **B:** 22 datasets from RNA fractions other than total mRNA. **C:** 2 datasets of clear cell renal carcinoma with VHL mutations and paired adjacent healthy tissues. **D:** 34 mouse RNA-seq datasets.

The next validation set consisted of four studies on specific fractions of RNA: newly transcribed RNA(4sU labeling RNA-seq and GRO-seq [38, 39]) and actively translated RNA (polysomal RNA-seq [40]) (Supplementary table S1-5, “RNA fractions”) [41–43]. In this case the challenge stems from the different RNA fractions used in the derivation of the trees (total mRNA) and test datasets. In spite of the different source of RNA, all decision trees had an accuracy over 0.75, with 36% showing an accuracy over 0.9 in the classification of 4sU labeling, GRO-seq, and polysomal samples (Fig. 2B).

The transcriptomic response to low oxygen tension can be induced by specific genetic lesions even under normoxia, so next we tested whether the classification trees could identify such samples, in spite of being derived from cells not exposed to hypoxia. Specifically we tested whether they could differentiate between clear cell renal carcinoma samples (ccRCC) and paired healthy adjacent tissues (Supplementary table S1-6, ”ccRCC”) from both the TCGA-KIRC collection and another publicly available study [44]. Over 80% of ccRCC show mutations in the von HippelLindau (VHL) gene that encodes for a key molecule controlling HIF stability. Thus VHL mutation leads to chronic HIF activation, even in the presence of oxygen, leading to a hypoxia-like transcriptional pattern [45, 46]. As shown in Fig. 2C, the vast majority of the trees, 216 out of 272, were able to identify VHL-mutant cells with an accuracy over 83%, even though no ccRCC samples were included in the training nor cross-validation datasets.

Finally, since the tree-classifiers were derived from human samples, we decided to test its performance on transcriptomes from other organisms. To that end, we gathered five studies in murine cells, totalling 34 individual RNA-seq experiments performed in different cell types and experimental conditions (Supplementary table S1-1 “RNA-seq Metadata”) [47–51]. As in the previous cases, the majority of trees (160 out 276) were able to classify samples with an accuracy of 79% or higher (2D). Altogether these results indicate that the tree-classifiers show a remarkable performance on novel datasets not used during the generation nor training steps, and correctly identify hypoxic samples derived from a wide range of conditions outside those represented in the training set. In spite of this, the bimodal distribution observed in Fig. 2C and 2D, suggest that a subset of the trees did not behave well on specific datasets. Closer analysis of these cases revealed that of the 58 trees showing poor performance against the ccRCC datasets, 42 share a common structure that includes only three genes organized in two levels, with ANKRD37 being the root nodes and two branches, one evaluating NDRG1 and the other BHLHE40 (Supplementary figure S1A). Although the expression of ANKRD37 differs in both groups (Supplementary figure S1B), both NDRG1 and BHLHE40 are already highly expressed in normal kidney samples (Supplementary figure S1C and D), explaining why trees with this topology are unable to differentiate between conditions. In the case of mouse datasets, we found that most of the best classifiers did not included MIR210HG in their structure (Fig. 2D), which stands to reason as there are no MIR210H orthologs annotated in mouse, therefore this feature does not convey relevant information for the classification in this case.

Altogether the analyses presented above allowed us to identify a subset of trees that accurately classify samples even from challenging datasets. Fig. 3A shows the best performing tree overall, especially apt in detecting short exposure to hypoxia and mildly low oxygen levels without overestimating the number of hypoxic samples in other validation sets. Since specific RNA types such as lncRNAs and microRNAs might not be represented in all sequencing libraries, we also selected the tree in Fig. 3B, being the best performing among those that don’t include MIR210HG lncRNA gene. Even though both trees perform reasonably well on mouse data, we have selected the additional tree in Fig. 3C for being the best performing in classifying murine samples specifically.

**Figure 3.**
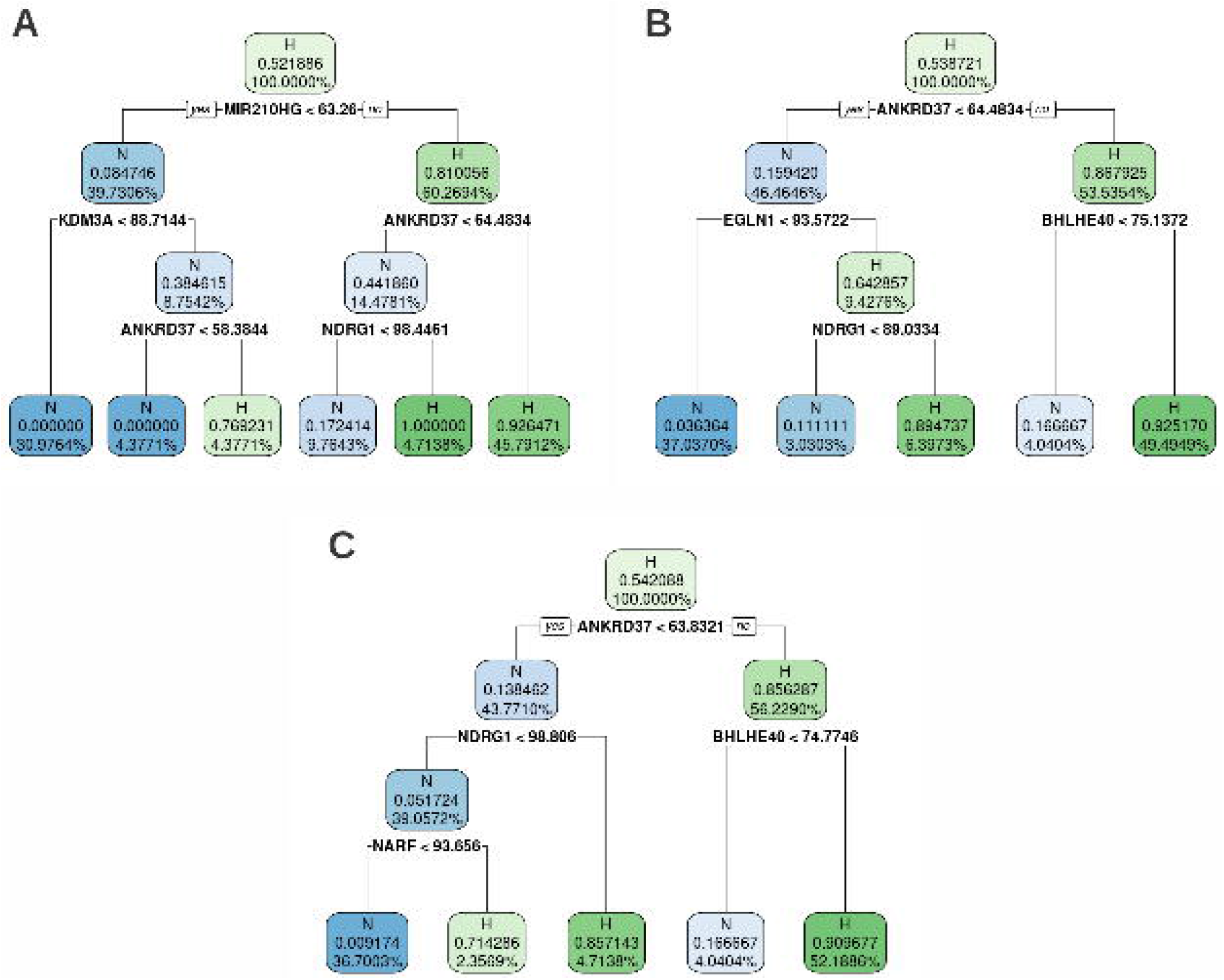
Selected decision trees. The labels in each node indicate the the node’s class (N, normoxic, and H, hypoxic), the probability of samples in the node to be classified as hypoxic, and the percentage of samples of the training set in each node. **A**: Overall best performing tree (tree #125). **B**: Best performing tree among those not using MIR210HG (tree #241) **C**: Best performing tree on mouse data (tree #42)

Since the selected trees derive from manual curation against a limited set of conditions outside those in the original datasets and it is unlikely that a single tree could accurately classify samples from all potential datasets, we tested whether classification would improve using several trees and generating a consensus. To this end, we compared the performance of the individual three selected trees (Fig. 4A-C) against three ensembles: the whole 276 collection (Fig. 4D), the 20 trees with higher mean F_1_-score (Fig. 4E) and the aforementioned three selected trees (Fig. 4F). AUC values for each curve are displayed in Table 1. The result of this analysis shows that, as expected, the ensembles tend to outperform individual trees. However, the difference is small and in some datasets individual datasets performed as well as the ensembles.

**Table 1.** Area under the curve corresponding to ROC curves for individual trees and three ensembles using four validation datasets. A sample was classified as hypoxic when the mean probability given by the ensemble or individual tree surpassed a given threshold between 0 and 1.

| Model | Time Series | RNA fractions | ccRC | Mouse |
| --- | --- | --- | --- | --- |
| Ensemble 276 Trees | 0.978 | 0.967 | 0.969 | 0.953 |
| Ensemble Top 20 | 0.991 | 1.000 | 0.990 | 0.946 |
| Ensemble selected 3 | 0.958 | 0.955 | 0.990 | 0.936 |
| Tree 125 | 0.958 | 0.955 | 0.944 | 0.910 |
| Tree 241 | 0.903 | 0.909 | 0.899 | 0.794 |
| Tree 42 | 0.903 | 0.909 | 0.656 | 0.882 |

**Figure 4.**
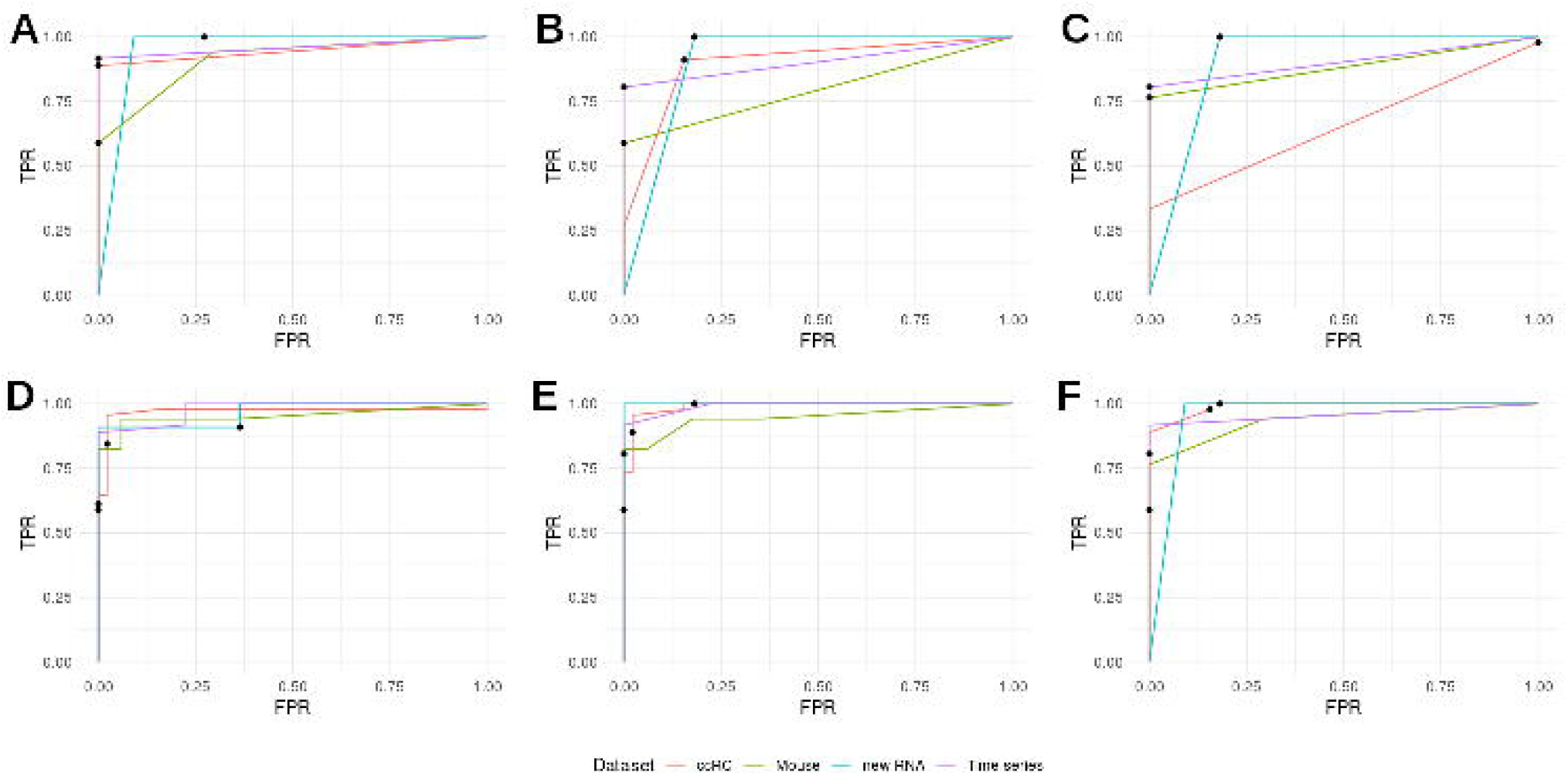
Tree ensembles performance. ROC curves for individual trees and three ensembles using four validation datasets. A sample was classified as hypoxic when the mean probability given by the trees surpassed a given threshold between 0 and 1. Black dots represent TPR/FPR values for a probability threshold of 0.5 to classify a sample as hypoxic. **A-C**: ROC curves for the individual trees selected for their performance. **A**: Tree #125 (Fig. 3A). **B**: Tree #241 (Fig. 3B). **C**: Tree #42 (Fig. 3C). **D-E**: ROC curves for three tree ensembles. **D**: all 276 trees. **E**: top 20 trees by medium F_1_-score. **F**: three selected trees (#125, #241, #42).

Thus, we have constructed a classification tree (tree #125, Fig. 3A,) that based on the ranked expression of just four genes (MIR210HG, KDM3A, ANKRD37 and NDRG1) is able to correctly identify normoxic/hypoxic samples with an accuracy of over 95% and 0.96 F_1_- score. The robustness of the predictions can be further improved by using the consensus decision from more than one tree.

### Application of the classifier to identify hypoxic tumors and hypoxic regions within tissues

Most solid tumors are hypoxic due to their aberrant growth and vascularization. Given that the presence of hypoxia compromises cancer therapy and is a poor prognosis factor, the identification of hypoxic tumors is relevant to predict tumor progression and select appropriate treatment strategies [52]. Thus, we next studied the ability of the tree classifier to identify hypoxic tumors. To that end we applied the classifier to transcriptomic profiles from the from The Cancer Genome Atlas (TCGA) and determined, for each type of tumor, the proportion of samples classified as hypoxic. As shown in Fig. 5A, the tumor type with the highest proportion of cases classified as hypoxic is the Kidney Renal Clear Cell Carcinoma (KIRC), in agreement with the molecular alterations characteristic of this cancer. Moreover, although the the ranking of tumors varies across studies [52, 53], head and neck and cervix carcinomas tend to be very hypoxic tumors as determined by direct measure of pO_2_ using oxygen electrodes, also in agreement with the classifier prediction shown in Fig. 5A.

**Figure 5.**
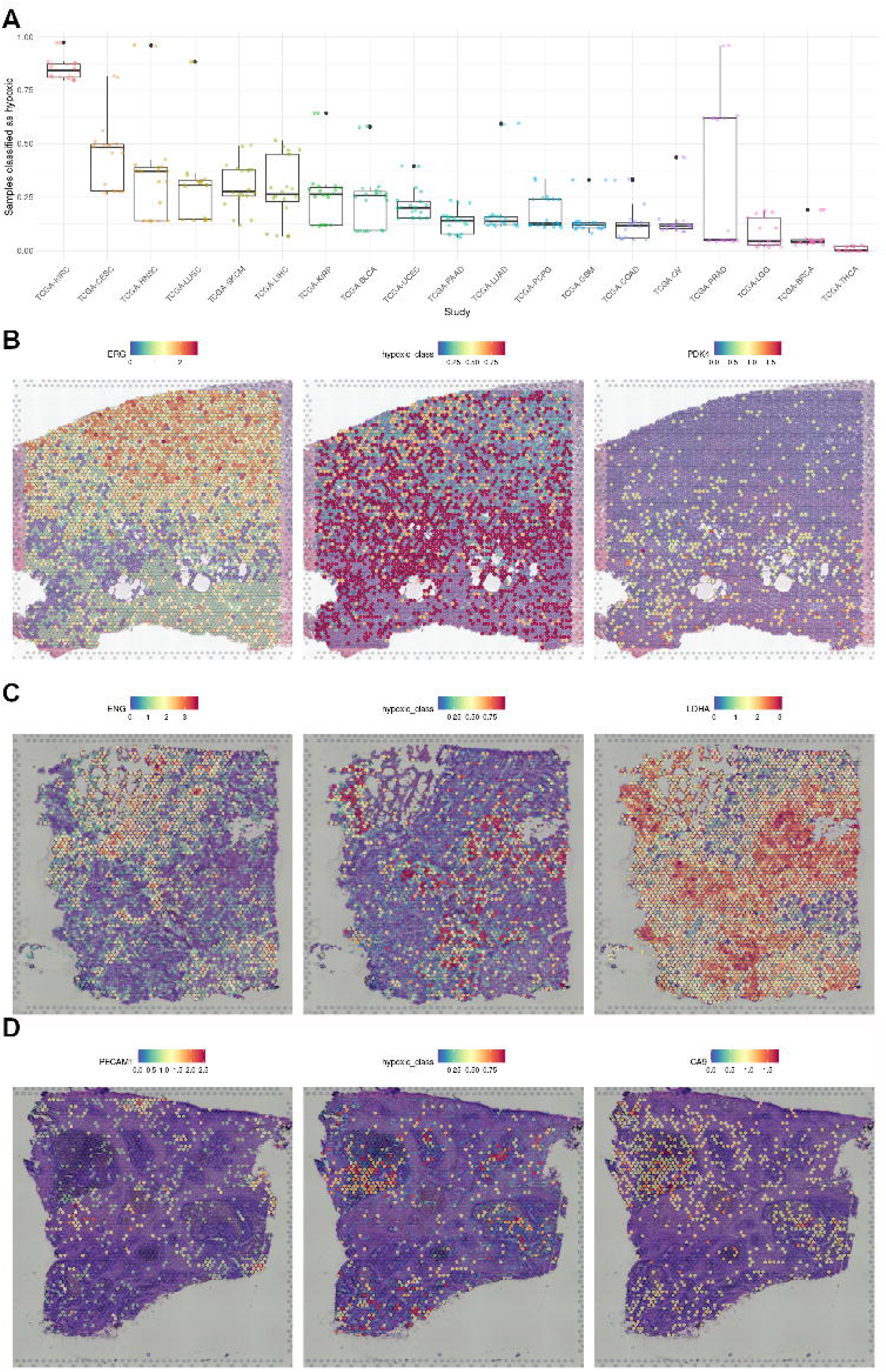
Detection of hypoxia in TCGA transcriptomes and tumor sections. **A**: Proportion of TCGA tumor samples classified as hypoxic by an ensemble of the 20 decision trees with higher F_1_-score, by primary site. **B-D**: Spatial Gene Expression datasets were downloaded from from 10X Genomics and used as input for the tree classifiers to detect hypoxic regions. On the left column, the expression of ENG, and PECAM1 are highlighted as endothelial markers, while on the right the expression of PDK4, LDHA, and CA9 mark regions of active anaerobic glycolysis. The central column represents the probability of each spot to be classified as hypoxic. **B**: Human Prostate Cancer, Adenocarcinoma with Invasive Carcinoma (FFPE). **C**: Human Glioblastoma. **D**: Human Colorectal Cancer.

One of the challenges in the study of tumor hypoxia is the heterogeneity of oxygenation within the tumoral mass [36]. The identification of hypoxic areas within a tumor typically relies on the detection of a single or a few markers of hypoxia such as the presence of HIFs or HIF targets [52]. The availability of spatial transcriptomic datasets allows for the identification of tissue hypoxia based on a gene signature rather than a single marker, so we decided to take advantage of the availability of several spatially resolved tumor sample transcriptomes [15, 17, 18] to test the ability of the tree classifiers to identify hypoxic regions in glioblastoma, prostate, and colorectal cancer. Each spot in the samples was classified as normoxic/hypoxic applying an ensemble of the 20 trees with higher mean F_1_-scores across validation datasets. For datasets that did not include MIR210HG expression, we generated an ensemble with trees that do not require this gene’s expression value.

This analysis revealed that regions identified as hypoxic by the tree classifiers, correspond to those poorly vascularized, according to vascular markers, and expressing high levels of glycolytic enzymes (Fig. 5B-D). It is worth mentioning that none of these reference markers were previously used to evaluate the performance of the classifiers.

In sharp contrast to the pervasive presence of hypoxic areas in most tumors, normal tissues usually do not show detectable HIF activity [54]. In order to test the specificity of the hypoxic signal detected by our classifiers, we next analyzed the spatial transcriptomes of normal tissues [19–21]. As shown in Fig. 6, with the exception of the kidney cortex, none of the normoxic tissues presented defined normoxic areas. These results are in agreement with a report showing that the kidneys are the only organ in showing HIF activity in normal mice breathing room air [55]. As further confirmation we proceeded to cluster the spots in each dataset (Supplementary figure S2) and examined differential expression between clusters containing high or low proportion of spots classified as hypoxic (Supplementary table S2-9 and Supplementary figure S3).

**Figure 6.**
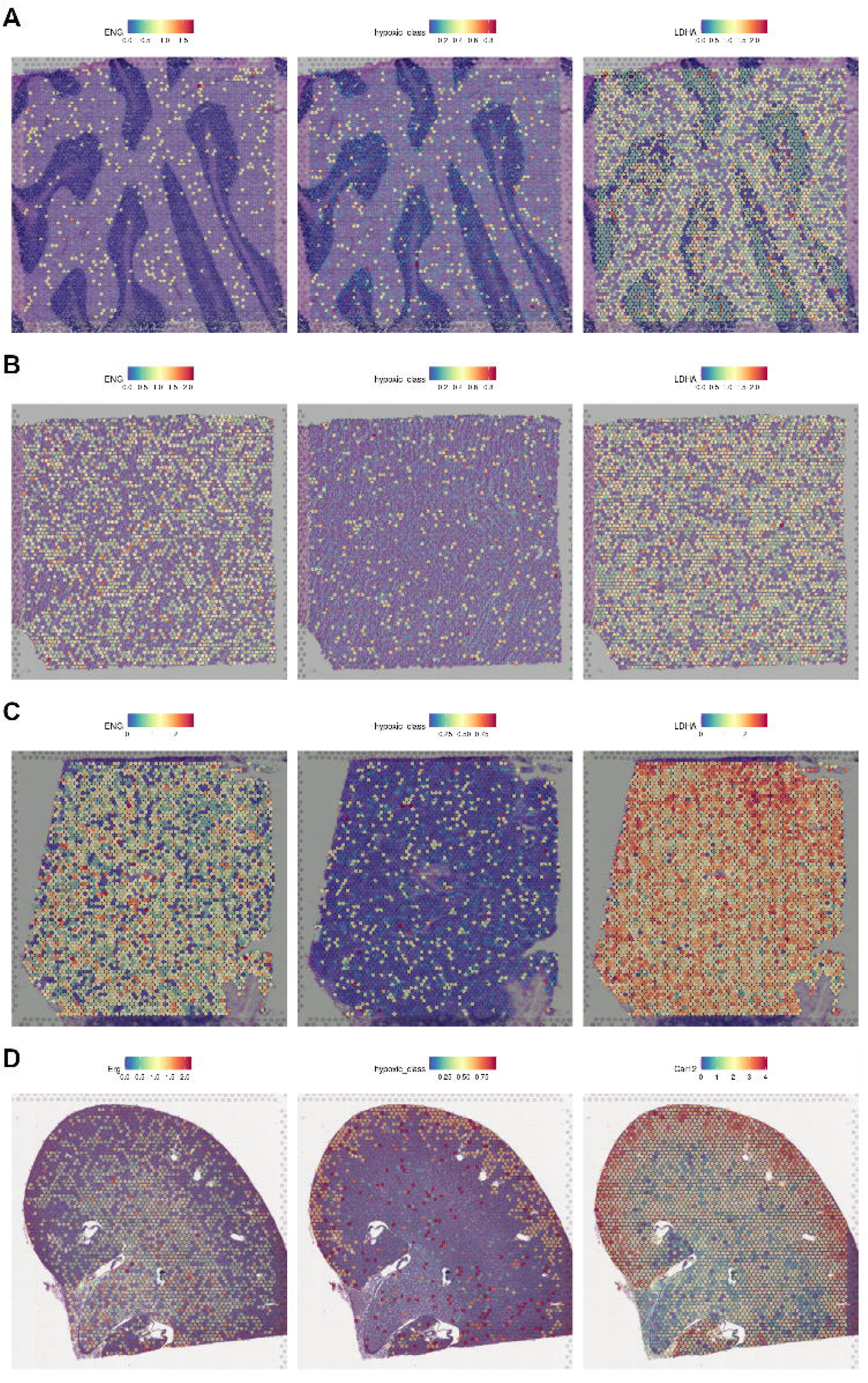
Detection of hypoxic regions within normal tissue sections.. Spatial Gene Expression datasets were downloaded from from 10X Genomics and used as input for the tree classifiers to detect hypoxic regions. On the left column, the expression of ERG, ENG are highlighted as endothelial markers, while on the right the expression of Car12, LDHA mark regions of active anaerobic glycolysis. The central column represents the probability of each spot to be classified as hypoxic.**A**: Cerebellum. **B**: Heart. **C**: Lymph node. **D**: Kidney.

Altogether, these results support the utility of our classifiers beyond bulk RNA-seq datasets, considering they accurately identify hypoxic tumoral samples and hypoxic regions in spatial gene expression datasets.

### Comparison with previously published hypoxia gene signatures

As indicated before, a number of hypoxic gene signatures have been previously described [2, 5–11], most of them derived from the lists of DEGs in response to hypoxia in specific tumors. Although these signatures are mostly defined as mere lists of genes and, as such, cannot be used to classify samples, Bhandari and coworkers ([56]) described a method to derive an hypoxic score value based on these lists of genes. Unlike the tree classifiers described herein, this score can not identify a sample as being hypoxic or normoxic, however, it allows allows the relative comparison among samples. We made use of this hypoxic scoring method to assess the relative ability of the individual gene signatures to discriminate between normoxic and hypoxic samples in the validation datasets described above (time series, RNA fractions, ccRCC and mouse RNA-seq datasets). Fig. 7A shows that the performance of the different gene signatures on the time series dataset varies widely and that only the scoring based on the Sorensen signature ([9]) results in a relative separation of samples that resembles their true labels. In the case of the different RNA fractions datasets, all gene signatures perform poorly as demonstrated by the very similar distribution of hypoxic scores assigned to normoxic and hypoxic samples (Fig. 7B), with only around 60% of the hypoxic samples having a score above those assigned to normoxic samples in the best cases. In contrast to these results, most signatures resulted in a good relative classification of normal and tumoral samples from the ccRCC datasets, as indicated by the score of the tumoral samples being higher than that of normal kidney ones (Fig. 7C). In spite of this, there was a substantial overlap between the two groups of samples for some signatures (Winter, Elvidge and Seigneuric2). Finally, we tested the gene signatures against samples from mouse cell lines, and as shown in Fig. 7D, even the best performing signatures (Sorensen and Elvidge), were unable to assign an score above controls to the majority of the hypoxic samples. Next, to directly compare the performance of the tree classifiers with the aforementioned gene signatures, we represented the hypoxic score assigned by each gene signature against the probability assigned by the ensemble of the 20 best trees for all the samples included in the validation datasets (time-course, RNA fractions, ccRCC and mouse RNA-seq samples). Fig. 7E shows that, although the two measures correlate for most gene signatures, the tree-based classifier described herein outperforms all gene signatures as evidenced by the better separation of samples according to the x-axis than the y-axis. Finally, Figs. 7F-G compare the performance of individual trees and gene signatures against each validation dataset. Remarkably, in the case of the most favorable dataset (clear cell renal carcinoma), individual trees perform similarly to the best gene signatures while thoroughly outperforming them in the rest of validation datasets.

**Figure 7.**
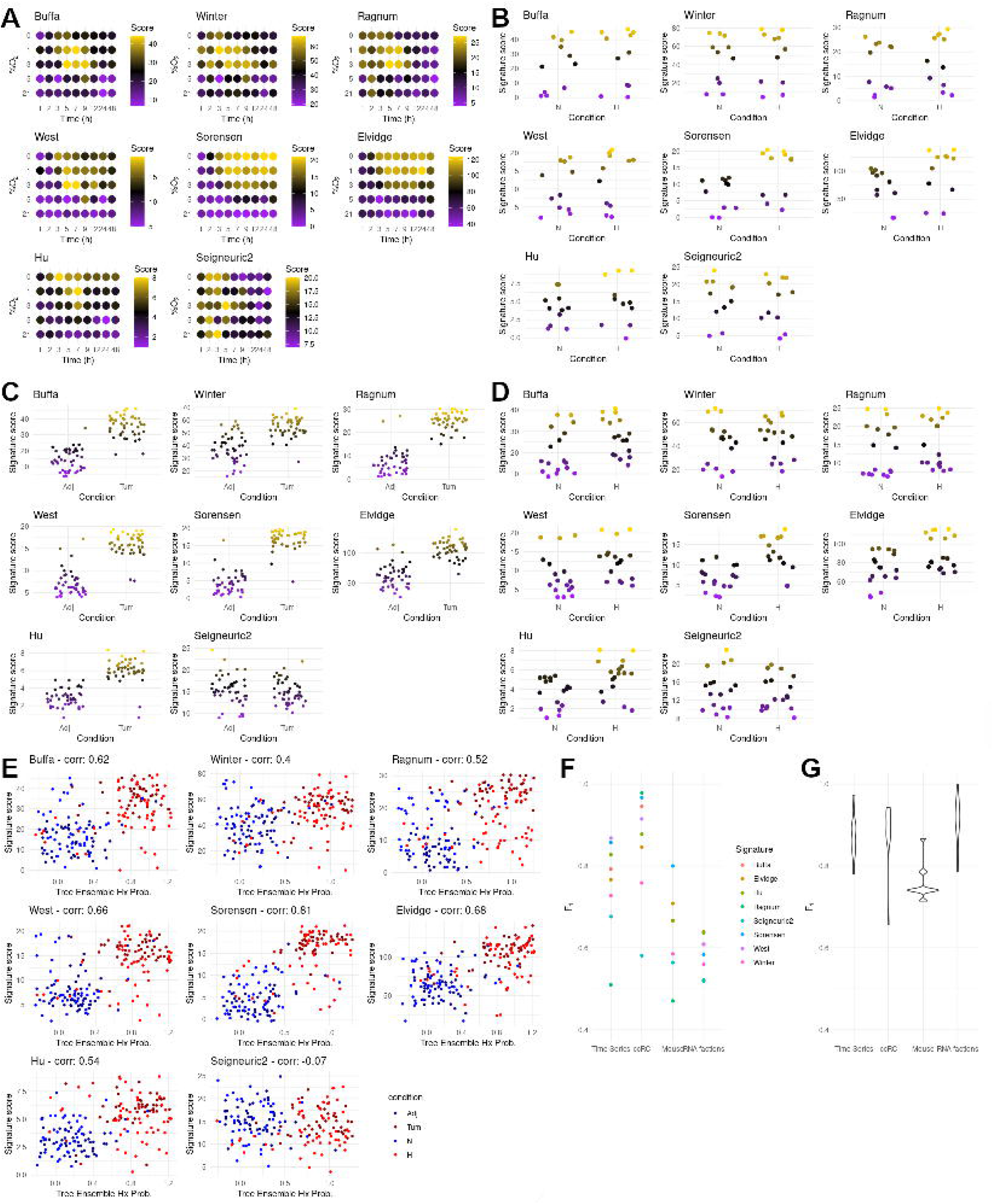
Performance of published hypoxic gene signatures. **A-D:** Application of 8 hypoxia gene signatures as described in [56] to training and validation datasets. **A**: PRJNA561635 time series. **B:** Specific RNA fractions other than total mRNA. **C**: ccRCC tumor and healthy adjacent tissue samples. **D**: mouse datasets. **E:** Correlation between the hypoxia scores derived from the 8 molecular signatures tested and the hypoxia probability calculated using an ensemble of our 20 best classifiers calculated for all samples from the validation experiments. Color code goes as follows; blue: samples grown in normoxia, red: samples grown in hypoxia, dark blue: healthy kidney samples, dark red: ccRC tumoral samples. **F-G:** distribution of F_1_-scores for normal and tumoral samples from the ccRCC dataset using the published gene signatures. **F:** 20 individual trees that composed the ensemble. Samples were classified as hypoxic when the probability given by a tree exceeded 0.5. **G:** Molecular signatures used in [56]. Samples were classified as hypoxic when the score calculated exceeded 50% of the maximum for that dataset and signature.

As a whole these results indicate that, in contrast to our classifiers, most of the published hypoxic gene signatures are less reliable when identifying cells exposed to hypoxia outside of the biological context each signatures was developed in. Basing our classifiers on the results of an extensive meta-analysis grants them the degree of flexibility needed to maintain accuracy against new data and different biological contexts.

## Discussion

In this work we aim to derive a gene signature that, besides defining the minimum core of genes that characterize the response to hypoxia, could be used to of assess if an individual gene expression dataset corresponds to sample that has been exposed to low oxygen tension. Additionally, one of the main priorities in the design of this classifier was to keep maximum transparency and interpretability in the process, so that, with a minimal or no background in machine learning, any user can not only determine if their sample is hypoxic, but also trace why it was marked as hypoxic. This work is based on a meta-analysis of the transcriptomic response to hypoxia, generated through the integration of a corpus of 69 differential expression datasets which included 425 individual RNA-seq experiments from 33 different cell types exposed to different degrees of hypoxia (0.1-5%O_2_) for a period of time spanning between 2 and 48h [24]. As a first filter of the variables (genes) to be included in the signature, we selected those widely expressed and significantly up-regulated by hypoxia according to the meta-analysis. This step ensured that the resulting models can be applied to a large variety of tissues as well as minimizing the risks of a biased corpus of publicly available experiments. Then we applied data mining methods to identify sets of genes that best separated normoxic and hypoxic samples using a treelike decision structure. Although the total number trees that achieved high accuracy was relatively large, only 16 out of the 20 pre-selected genes were required among all the trees, with many having the same structure and differing only slightly in the gene expression threshold. Moreover, the vast majority of trees included different combination of 3-5 genes from the set EGLN1, MIR210HG, NDRG1, ANKRD37, TCAF2, PFKFB3, BHLHE40, and MAFF (Supplementary table S1).

In contrast with classical molecular signatures, the trees described herein provide not just a list of genes relevant to the process, but also a set of matching quantitative expression boundaries, which allows it to classify individual samples both from a binary perspective (hypoxic or normoxic sample) as well as a continuous one (probability of a sample to be classified as hypoxic, shown in Fig. 5B-D and 6). The features of the classifiers permit their application of the classification trees to a wide range of gene expression datasets, from the conventional bulk RNA-seq by polyA capture and techniques to characterize newly transcribed RNA [38, 39] to spatially resolved transcriptomics and single cell RNA-seq. Importantly, gene’s expression boundaries are represented as the percentile occupied by the gene in a ranked list of expression values from a given sample, which means that this method can be applied to a diverse set of input formats: raw reads, counts per million, FPKM, variance stabilizing transformations, etc.

It should be noted that these classifiers are robust enough to predict the condition of samples from murine cells despite having being trained only with human datasets, as well as identify samples in which response to hypoxia is activated by mutations in specific signalling pathways (ccRCC dataset), due to the pattern of vascularization and/or oxygen consumption (TCGA datasets, tumor specimens) and even hypoxic regions present in normal tissues (kidney dataset).

In regard to the classification of tumor samples according to their degree of hypoxia, our results are in good agreement to those reported in [56] using different hypoxic signatures (Supplementary table S1-7). However, unlike the tree classifiers described herein, other signatures failed to identify clear cell renal carcinomas as the type of tumor showing the highest up-regulation of the hypoxic transcriptome [56]. Nevertheless, as shown in Fig. 5A, with the exception of renal carcinomas, the proportion of tumor samples classified as hypoxic in each group resembles more closely the results obtained by Bhandari et al. ([56]). On the other hand, previously reported hypoxic signatures performed poorly against non-tumoral validation datasets described in our work, as shown in Fig. 7 and supplementary table S3. Considering that all but one of the classic gene signatures were defined in the context of tumor hypoxia, a poor correlation could be expected when compared to the performance of a classifier trained with a more diverse corpus of experiments (Fig. 7E).

As further confirmation of the effectiveness of the trees in comparison to classic gene signatures, we decided to test them in spatial gene expression datasets, using the expression levels of endothelial markers ERG, ENG and PECAM1/CD31 to localize well oxygenated regions and the expression levels of genes related to anaerobic glycolysis, to define regions of restricted oxygen availability. As shown in Fig. 5B-D, regions classified as hypoxic overlap those of active anaerobic glycolysis, meanwhile regions rich in endothelial markers tend to be classified as normoxic. After unsupervised clustering of the same datasets (Fig. S2), differential expression between clusters overlapping normoxic and hypoxic areas highlighted genes linked to hypoxia and not included in our models, such as VEGFA or ENO1 (Supplementary table S2). Furthermore, when comparing the adjusted p-values of genes up-regulated between clusters that are also significantly up-regulated in the cited hypoxia meta-analysis [24] (random effect>0.7 and FDR<0.01) this group has significantly lower p-values than genes not linked to hypoxia, confirming an enrichment on hypoxia-related genes among those differentially expressed between hypoxic and normoxic clusters. In contrast to the results obtained with tumor sections, we did not found significant hypoxic regions in normal tissues (Fig. 6). Which is consistent with the absence of HIF activation in tissues under physiological conditions [54], in spite the wide range of pO2 values found in normal tissues [36]. The only exception was the identification of the kidney cortex as an hypoxic region (Fig. 6), which, although unexpected at first glance, is in agreement with the results from a noninvasive imaging technique that identified the kidneys as the only organ in showing HIF activity in normal mice breathing room air [55]. Moreover, although pO_2_ in the medulla is lower than in the cortex, the renal medulla presents a comparatively higher expression of the HIF inhibitors EGLNs [57], which might explain why no constitutive HIF stabilization is found in the medulla under physiological conditions [58] and thus why this region is not labeled by the tree classifier.

In addition to their remarkable performance, the structure of the decision trees allows for biological interpretation of the prediction’s results. In this regard, the application of the decision trees to the challenging datasets provided relevant and novel insights into the underlying biological processes. For example, the analysis of the performance of different trees on the ccRCC dataset revealed that BHLHE40 and NDRG1 are expressed at high levels in renal tissue which can hint to specific functions of these genes in kidney physiology. On the other hand, as seen with the mouse dataset in Fig. 2D, missing data in one of the classifying variables (MIR210HG) could directly or indirectly hinder the performance of the trees. Thus, we tested if performance of the tree classifiers can be improved by generating a consensus. As we show in Fig. 4, an ensemble of the 20 trees with higher mean F_1_-score (Fig. 4D) can outperform all individual trees and other ensembles in most cases (with the exception of tree #42 in the mouse dataset). A classification based just in the consensus of the three trees selected in this paper (Fig. 4F) can compensate for the shortcomings of each individual model while maintaining the ease of use intended for this classifier. Tree ensembles could be a better suited alternative for samples that are harder to classify or derived from a dataset distantly related to the ones used to derive our tree classifiers.

In summary, herein we describe a ensemble of tree gene signatures that can be easily implemented to identify hypoxic samples based on their transcriptomic profile without the need for a reference. Given the importance of oxygen homeostasis in physiology and disease, this tool could be useful in a wide variety of research and clinical settings. Finally, in the view of its merits, we proposed the extension of this method to define gene signatures that characterize other cellular processes.

## Declarations

### Ethics approval and consent to participate

Not applicable

### Consent for publication

Not applicable

### Availability of data and materials

References to the sources of experiments used as validation datasets are included in Supplementary table S1, sheet 1. The full collection of 276 decision trees, gene ranking percentiles for the validation datasets used, as well as a step-by-step guide to apply the classifiers to new data are available at the following GitHub repository: www.github.com/LauraPS1/Hypoxia_Classifier.

### Authors’ contributions

- **LPS**: Downloading and processing raw data of 220 of the 425 RNA-seq samples used to generate the classifiers, as well as the bulk and spatial RNA-seq experiments used as validation datasets. Methodology development, generation and validation of the classifiers, and manuscript writing, review and editing.
- **LSG**: Downloading and processing raw data of 205 of the 425 RNA-seq samples used to generate the classifiers
- **RRR**: Acquisition of the financial support for the project leading to this publication, advise and revision during the writing process.
- **LP**: Conceptualization and project administration, methodology development, and manuscript writing, review and editing.

## Acknowledgements

The results shown here are in part based upon data generated by the TCGA Research Network: www.cancer.gov/tcga.

---

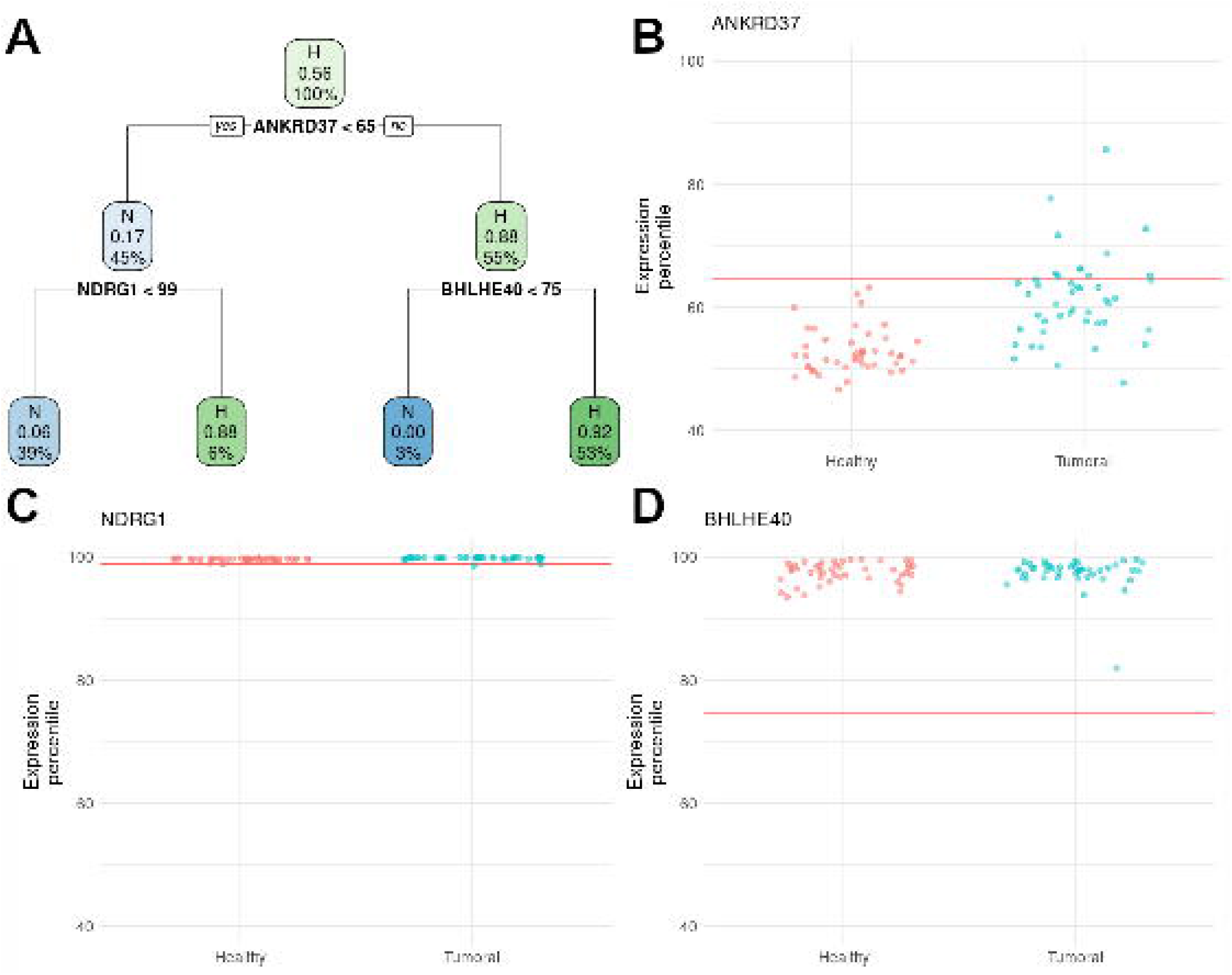

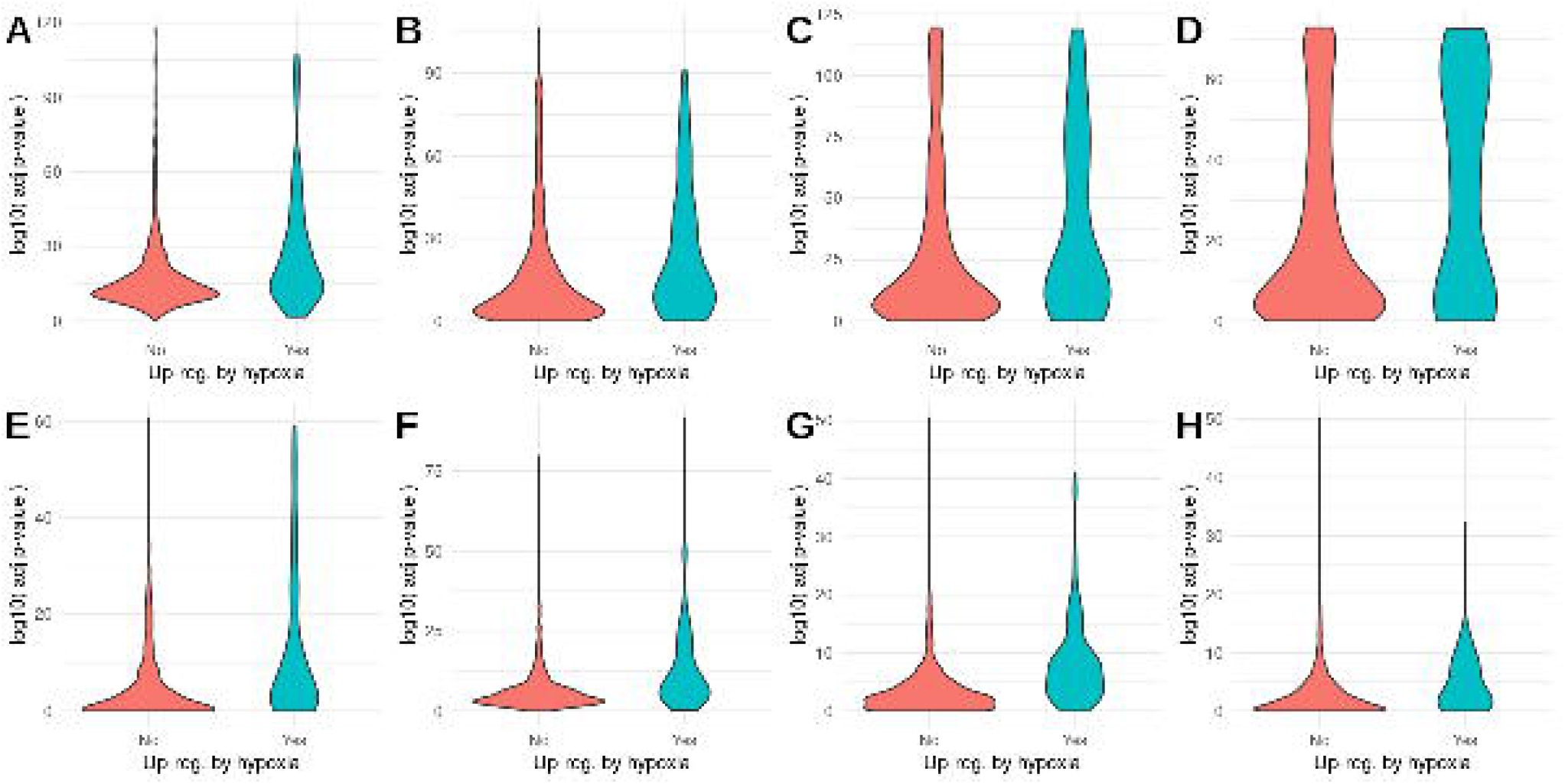

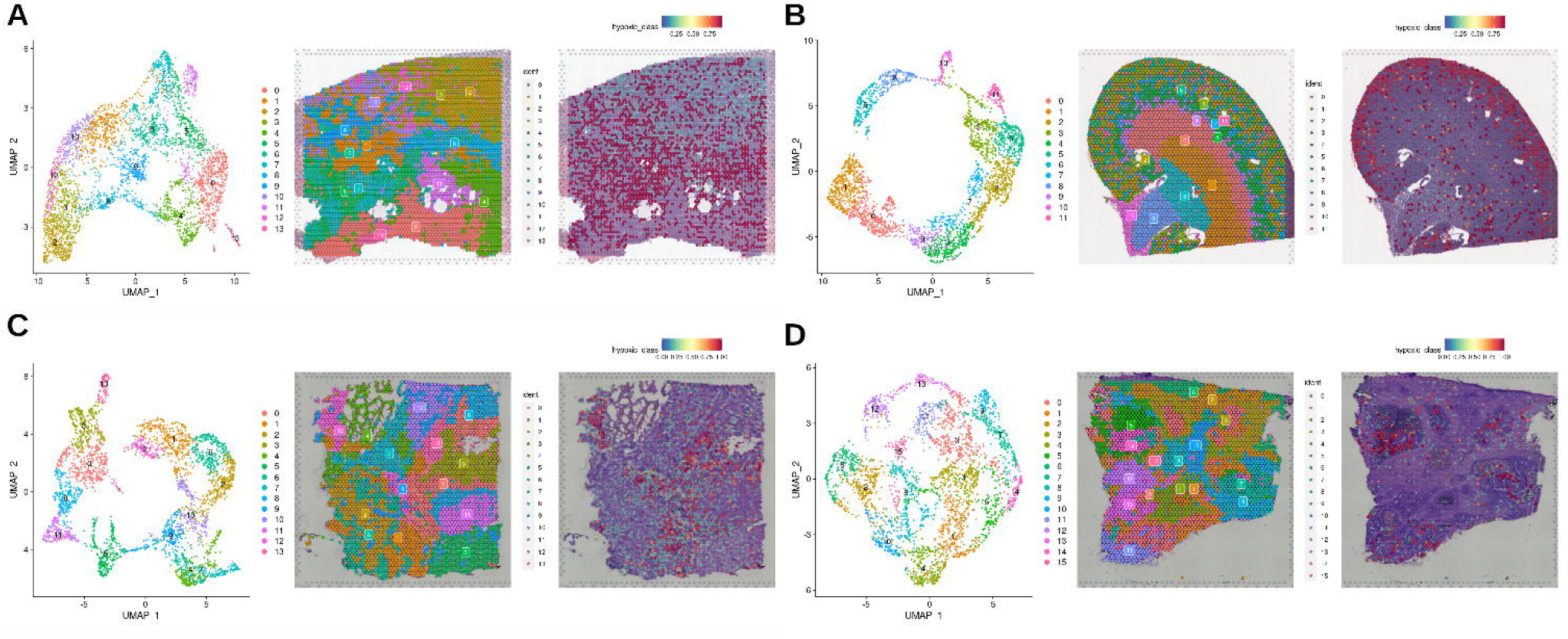

## References

1. Semenza, G.L.: Hypoxia-inducible factors in physiology and medicine. Cell 148(3), 399–408 (2012)

2. Elvidge, G.P., Glenny, L., Appelhoff, R.J., Ratcliffe, P.J., Ragoussis, J., Gleadle, J.M.: Concordant regulation of gene expression by hypoxia and 2-oxoglutarate-dependent dioxygenase inhibition: The role of hif-1α, hif-2α, and other pathways*. Journal of Biological Chemistry 281(22), 15215–15226 (2006). doi:10.1074/jbc.M511408200

3. Yang, G., Shi, R., Zhang, Q.: Hypoxia and oxygen-sensing signaling in gene regulation and cancer progression. International Journal of Molecular Sciences 21(21) (2020). doi:10.3390/ijms21218162

4. Mesarwi, O.A., Loomba, R., Malhotra, A.: Obstructive sleep apnea, hypoxia, and nonalcoholic fatty liver disease. American Journal of Respiratory and Critical Care Medicine 199(7), 830–841 (2019). doi:10.1164/rccm.201806-1109TR. https://doi.org/10.1164/rccm.201806-1109TR

5. Buffa, F.M., Harris, A.L., West, C.M., Miller, C.J.: Large meta-analysis of multiple cancers reveals a common, compact and highly prognostic hypoxia metagene. British Journal of Cancer 102(2), 428–435 (2010). doi:10.1038/sj.bjc.6605450

6. Winter, S.C., Buffa, F.M., Silva, P., Miller, C., Valentine, H.R., Turley, H., Shah, K.A., Cox, G.J., Corbridge, R.J., Homer, J.J., Musgrove, B., Slevin, N., Sloan, P., Price, P., West, C.M.L., Harris, A.L.: Relation of a hypoxia metagene derived from head and neck cancer to prognosis of multiple cancers. Cancer Research 67(7), 3441–3449 (2007). doi:10.1158/0008-5472.CAN-06-3322. https://cancerres.aacrjournals.org/content/67/7/3441.full.pdf

7. Ragnum, H.B., Vlatkovic, L., Lie, A.K., Axcrona, K., Julin, C.H., Frikstad, K.M., Hole, K.H., Seierstad, T., Lyng, H.: The tumour hypoxia marker pimonidazole reflects a transcriptional programme associated with aggressive prostate cancer. British Journal of Cancer 112(2), 382–390 (2015). doi:10.1038/bjc.2014.604

8. Eustace, A., Mani, N., Span, P.N., Irlam, J.J., Taylor, J., Betts, G.N.J., Denley, H., Miller, C.J., Homer, J.J., Rojas, A.M., Hoskin, P.J., Buffa, F.M., Harris, A.L., Kaanders, J.H.A.M., West, C.M.L.: A 26-gene hypoxia signature predicts benefit from hypoxia-modifying therapy in laryngeal cancer but not bladder cancer. Clinical Cancer Research 19(17), 4879–4888 (2013). doi:10.1158/1078-0432.CCR-13-0542. https://clincancerres.aacrjournals.org/content/19/17/4879.full.pdf

9. Sørensen, B.S., Toustrup, K., Horsman, M.R., Overgaard, J., Alsner, J.: Identifying ph independent hypoxia induced genes in human squamous cell carcinomas in vitro. Acta Oncologica 49(7), 895–905 (2010). doi:10.3109/02841861003614343. https://doi.org/10.3109/02841861003614343

10. Hu, Z., Fan, C., Livasy, C., He, X., Oh, D.S., Ewend, M.G., Carey, L.A., Subramanian, S., West, R., Ikpatt, F., Olopade, O.I., van de Rijn, M., Perou, C.M.: A compact VEGF signature associated with distant metastases and poor outcomes. BMC Medicine 7 (2009). doi:10.1186/1741-7015-7-9

11. Seigneuric, R., Starmans, M.H.W., Fung, G., Krishnapuram, B., Nuyten, D.S.A., van Erk, A., Magagnin, M.G., Rouschop, K.M., Krishnan, S., Rao, R.B., Evelo, C.T.A., Begg, A.C., Wouters, B.G., Lambin, P.: Impact of supervised gene signatures of early hypoxia on patient survival. Radiotherapy and Oncology 83(3), 374–382 (2007). doi:10.1016/j.radonc.2007.05.002

12. Leinonen, R., Sugawara, H., Shumway, M.: The sequence read archive. Nucleic Acids Research 39(SUPPL. 1), 148–162 (2011). doi:10.1093/nar/gkq1019

13. Patro, R., Duggal, G., Love, M.I., Irizarry, R.A., Kingsford, C.: Salmon provides fast and bias-aware quantification of transcript expression. Nature Methods 14(4), 417–419 (2017). doi:10.1038/nmeth.4197

14. O’Leary, N.A., Wright, M.W., Brister, J.R., Ciufo, S., Haddad, D., McVeigh, R., Rajput, B., Robbertse, B., Smith-White, B., Ako-Adjei, D., Astashyn, A., Badretdin, A., Bao, Y., Blinkova, O., Brover, V., Chetvernin, V., Choi, J., Cox, E., Ermolaeva, O., Farrell, C.M., Goldfarb, T., Gupta, T., Haft, D., Hatcher, E., Hlavina, W., Joardar, V.S., Kodali, V.K., Li, W., Maglott, D., Masterson, P., McGarvey, K.M., Murphy, M.R., O’Neill, K., Pujar, S., Rangwala, S.H., Rausch, D., Riddick, L.D., Schoch, C., Shkeda, A., Storz, S.S., Sun, H., Thibaud-Nissen, F., Tolstoy, I., Tully, R.E., Vatsan, A.R., Wallin, C., Webb, D., Wu, W., Landrum, M.J., Kimchi, A., Tatusova, T., DiCuccio, M., Kitts, P., Murphy, T.D., Pruitt, K.D.: Reference sequence (RefSeq) database at NCBI: Current status, taxonomic expansion, and functional annotation. Nucleic Acids Research 44(D1), 733–745 (2016). doi:10.1093/nar/gkv1189

15. Genomics, X.: Human Prostate Cancer, Adenocarcinoma with Invasive Carcinoma (FFPE), Spatial Gene Expression Dataset by Space Ranger 1.3.0 (2021)

16. Genomics, X.: Adult Mouse Kidney (FFPE), Spatial Gene Expression Dataset by Space Ranger 1.3.0 (2021)

17. Genomics, X.: Human Colorectal Cancer: Whole Transcriptome Analysis, Spatial Gene Expression Dataset by Space Ranger 1.2.0 (2020)

18. Genomics, X.: Human Glioblastoma: Whole Transcriptome Analysis, Spatial Gene Expression Dataset by Space Ranger 1.2.0 (2020)

19. Genomics, X.: Human Cerebellum: Whole Transcriptome Analysis, Spatial Gene Expression Dataset by Space Ranger 1.2.0 (2020)

20. Genomics, X.: Human Heart, Spatial Gene Expression Dataset by Space Ranger 1.1.0 (2020)

21. Genomics, X.: Human Lymph Node, Spatial Gene Expression Dataset by Space Ranger 1.2.0 (2020)

22. Hafemeister, C., Satija, R.: Normalization and variance stabilization of single-cell RNA-seq data using regularized negative binomial regression. Genome Biology 20(1) (2019). doi:10.1186/s13059-019-1874-1

23. Hao, Y., Hao, S., Andersen-Nissen, E., III, W.M.M., Zheng S., Butler, A., Lee, M.J., Wilk, A.J., Darby, C., Zagar, M., Hoffman, P., Stoeckius, M., Papalexi, E., Mimitou, E.P., Jain, J., Srivastava, A., Stuart, T., Fleming, L.B., Yeung, B., Rogers, A.J., McElrath, J.M., Blish, C.A., Gottardo, R., Smibert, P., Satija, R.: Integrated analysis of multimodal single-cell data. Cell (2021). doi:10.1016/j.cell.2021.04.048

24. Puente-Santamaria, L., Sanchez-Gonzalez, L., Gonzalez-Serrano, B.P., Pescador, N., Martinez-Costa, O.H., Ramos-Ruiz, R., del Peso, L.: Formal meta-analysis of hypoxic gene expression profiles reveals a universal gene signature and cell type-specific effects. bioRxiv (2021). doi:10.1101/2021.11.12.468418. https://www.biorxiv.org/content/early/2021/11/14/2021.11.12.468418.full.pdf

25. Liaw, A., Wiener, M.: Classification and regression by randomforest. R News 2(3), 18–22 (2002)

26. Therneau, T., Atkinson, B.: Rpart: Recursive Partitioning and Regression Trees. (2019). R package version 4.1-15. https://CRAN.R-project.org/package=rpart

27. Kaelin, W.G., Ratcliffe, P.J.: Oxygen sensing by metazoans: The central role of the hif hydroxylase pathway. Molecular Cell 30(4), 393–402 (2008). doi:10.1016/j.molcel.2008.04.009

28. Huang, X., Le, Q.T., Giaccia, A.J.: MiR-210 - micromanager of the hypoxia pathway. Trends in Molecular Medicine 16(5), 230–237 (2010). doi:10.1016/j.molmed.2010.03.004

29. Cangul, H.: Hypoxia upregulates the expression of the NDRG1 gene leading to its overexpression in various human cancers. BMC Genetics 5 (2004). doi:10.1186/1471-2156-5-27

30. Benita, Y., Kikuchi, H., Smith, A.D., Zhang, M.Q., Chung, D.C., Xavier, R.J.: An integrative genomics approach identifies Hypoxia Inducible Factor-1 (HIF-1)-target genes that form the core response to hypoxia. Nucleic Acids Research 37(14), 4587–4602 (2009). doi:10.1093/nar/gkp425. https://academic.oup.com/nar/article-pdf/37/14/4587/16752655/gkp425.pdf

31. Saeki, K., Onishi, H., Koga, S., Ichimiya, S., Nakayama, K., Oyama, Y., Kawamoto, M., Sakihama, K., Yamamoto, T., Matsuda, R., Miyasaka, Y., Nakamura, M., Oda, Y.: Fam115c could be a novel tumor suppressor associated with prolonged survival in pancreatic cancer patients. J Cancer 11, 2289–2302 (2020). doi:10.7150/jca.38399

32. Obach, M., Àurea Navarro-Sabaté Caro, J., Kong, X., Duran, J., Gómez, M., Perales, J.C., Ventura, F., Rosa, J.L., Bartrons, R.: 6-phosphofructo-2-kinase (pfkfb3) gene promoter contains hypoxia-inducible factor-1 binding sites necessary for transactivation in response to hypoxia*. Journal of Biological Chemistry 279(51), 53562–53570 (2004). doi:10.1074/jbc.M406096200

33. Ivanova, A.V., Ivanov, S.V., Danilkovitch-Miagkova, A., Lerman, M.I.: Regulation of stra13 by the von hippel-lindau tumor suppressor protein, hypoxia, and the ubc9/ubiquitin proteasome degradation pathway*. Journal of Biological Chemistry 276(18), 15306–15315 (2001). doi:10.1074/jbc.M010516200

34. Chen, L., Fink, T., Ebbesen, P., Zachar, V.: Temporal transcriptome of mouse atdc5 chondroprogenitors differentiating under hypoxic conditions. Experimental Cell Research 312(10), 1727–1744 (2006). doi:10.1016/j.yexcr.2006.02.013

35. Klomp, J., Hyun, J., Klomp, J.E., Pajcini, K., Rehman, J., Malik, A.B.: Comprehensive transcriptomic profiling reveals sox7 as an early regulator of angiogenesis in hypoxic human endothelial cells. Journal of Biological Chemistry 295(15), 4796–4808 (2020). doi:10.1074/jbc.RA119.011822

36. McKeown, S.R.: Defining normoxia, physoxia and hypoxia in tumours - Implications for treatment response. British Journal of Radiology 87(1035) (2014). doi:10.1259/bjr.20130676

37. Löfstedt, T., Fredlund, E., Holmquist-Mengelbier, L., Pietras, A., Ovenberger, M., Poellinger, L., Påhlman, S.: Hypoxia inducible factor-2α in cancer. Cell Cycle 6(8), 919–926 (2007). doi:10.4161/cc.6.8.4133. PMID: 17404509. https://doi.org/10.4161/cc.6.8.4133

38. Garibaldi, A., Carranza, F., Hertel, K.J.: Isolation of newly transcribed rna using the metabolic label 4-thiouridine. Methods in Molecular Biology 1648, 169–176 (2017). doi:10.1007/978-1-4939-7204-313

39. Gardini, A.: Global run-on sequencing (GRO-Seq). Methods in Molecular Biology 1468, 111–120 (2017). doi:10.1007/978-1-4939-4035-69

40. Panda, A., Martindale, J., Gorospe, M.: Polysome Fractionation to Analyze mRNA Distribution Profiles. Bio-Protocol 7(3) (2017). doi:10.21769/bioprotoc.2126

41. Tiana, M., Acosta-Iborra, B., Puente-Santamaría, L., Hernansanz-Agustin, P., Worsley-Hunt, R., Masson, N., García-Rio, F., Mole, D., Ratcliffe, P., Wasserman, W.W., Jimenez, B., del Peso, L.: The SIN3A histone deacetylase complex is required for a complete transcriptional response to hypoxia. Nucleic Acids Research 46(1), 120–133 (2018). doi:10.1093/nar/gkx951

42. Niskanen, H., Tuszynska, I., Zaborowski, R., Heinäniemi, M., Ylä-Herttuala, S., Wilczynski, B., Kaikkonen, M.U.: Endothelial cell differentiation is encompassed by changes in long range interactions between inactive chromatin regions. Nucleic Acids Research 46(4), 1724–1740 (2018). doi:10.1093/nar/gkx1214

43. Sesé, M., Fuentes, P., Esteve-Codina, A., Béjar, E., McGrail, K., Thomas, G., Aasen, T., Ramón y Cajal, S.: Hypoxia-mediated translational activation of ITGB3 in breast cancer cells enhances TGF-β signaling and malignant features in vitro and in vivo. Oncotarget 8(70), 114856–114876 (2017). doi:10.18632/oncotarget.23145

44. Yao, X., Tan, J., Lim, K.J., Koh, J., Ooi, W.F., Li, Z., Huang, D., Xing, M., Chan, Y.S., Qu, J.Z., Tay, S.T., Wijaya, G., Lam, Y.N., Hong, J.H., Lee-Lim, A.P., Guan, P., Ng, M.S.W., He, C.Z., Lin, J.S., Nandi, T., Qamra, A., Xu, C., Myint, S.S., Davies, J.O.J., Goh, J.Y., Loh, G., Tan, B.C., Rozen, S.G., Yu, Q., Tan, I.B.H., Cheng, C.W.S., Li, S., Chang, K.T.E., Tan, P.H., Silver, D.L., Lezhava, A., Steger, G., Hughes, J.R., Teh, B.T., Tan, P.: VHL deficiency drives enhancer activation of oncogenes in clear cell renal cell carcinoma. Cancer Discovery 7(11), 1284–1305 (2017). doi:10.1158/2159-8290.CD-17-0375

45. Guo, G., Gui, Y., Gao, S., Tang, A., Hu, X., Huang, Y., Jia, W., Li, Z., He, M., Sun, L., Song, P., Sun, X., Zhao, X., Yang, S., Liang, C., Wan, S., Zhou, F., Chen, C., Zhu, J., Li, X., Jian, M., Zhou, L., Ye, R., Huang, P., Chen, J., Jiang, T., Liu, X., Wang, Y., Zou, J., Jiang, Z., Wu, R., Wu, S., Fan, F., Zhang, Z., Liu, L., Yang, R., Liu, X., Wu, H., Yin, W., Zhao, X., Liu, Y., Peng, H., Jiang, B., Feng, Q., Li, C., Xie, J., Lu, J., Kristiansen, K., Li, Y., Zhang, X., Li, S., Wang, J., Yang, H., Cai, Z., Wang, J.: Frequent mutations of genes encoding ubiquitin-mediated proteolysis pathway components in clear cell renal cell carcinoma. Nature Genetics 44(1), 17–19 (2012). doi:10.1038/ng.1014

46. Chittiboina, P., Lonser, R.R.: Chapter 10 - von hippel–lindau disease. In: Islam, M.P., Roach, E.S. (eds.) Neurocutaneous Syndromes. Handbook of Clinical Neurology, vol. 132, pp. 139–156. Elsevier, ??? (2015). doi:10.1016/B978-0-444-62702-5.00010-X. https://www.sciencedirect.com/science/article/pii/B978044462702500010X

47. Bischoff, F.C., Werner, A., John, D., Boeckel, J.N., Melissari, M.T., Grote, P., Glaser, S.F., Demolli, S., Uchida, S., Michalik, K.M., Meder, B., Katus, H.A., Haas, J., Chen, W., Pullamsetti, S.S., Seeger, W., Zeiher, A.M., Dimmeler, S., Zehendner, C.M.: Identification and Functional Characterization of Hypoxia-Induced Endoplasmic Reticulum Stress Regulating lncRNA (HypERlnc) in Pericytes. Circulation Research 121(4), 368–375 (2017). doi:10.1161/CIRCRESAHA.116.310531

48. MacVicar, T., Ohba, Y., Nolte, H., Mayer, F.C., Tatsuta, T., Sprenger, H.G., Lindner, B., Zhao, Y., Li, J., Bruns, C., Krüger, M., Habich, M., Riemer, J., Schwarzer, R., Pasparakis, M., Henschke, S., Brüning, J.C., Zamboni, N., Langer, T.: Lipid signalling drives proteolytic rewiring of mitochondria by YME1L. Nature 575(7782), 361–365 (2019). doi:10.1038/s41586-019-1738-6

49. Lo, K.A., Labadorf, A., Kennedy, N.J., Han, M.S., Yap, Y.S., Matthews, B., Xin, X., Sun, L., Davis, R.J., Lodish, H.F., Fraenkel, E.: Analysis of In Vitro Insulin-Resistance Models and Their Physiological Relevance to InVivo Diet-Induced Adipose Insulin Resistance. Cell Reports 5(1), 259–270 (2013). doi:10.1016/j.celrep.2013.08.039

50. Kiernan, E.A., Ewald, A.C., Ouellette, J.N., Wang, T., Roopra, A., Watters, J.J.: Prior hypoxia exposure enhances murine microglial inflammatory gene expression in vitro without concomitant H3K4me3 enrichment. bioRxiv (2020). doi:10.1101/2020.02.03.933028

51. Islam, M.R., Lbik, D., Sakib, M.S., Maximilian Hofmann, R., Berulava, T., Jiménez Mausbach, M., Cha, J., Goldberg, M., Vakhtang, E., Schiffmann, C., Zieseniss, A., Katschinski, D.M., Sananbenesi, F., Toischer, K., Fischer, A.: Epigenetic gene expression links heart failure to memory impairment. EMBO Molecular Medicine 13(3) (2021). doi:10.15252/emmm.201911900

52. Walsh, J.C., Lebedev, A., Aten, E., Madsen, K., Marciano, L., Kolb, H.C.: The clinical importance of assessing tumor hypoxia: relationship of tumor hypoxia to prognosis and therapeutic opportunities. Antioxid Redox Signal 21(10), 1516–1554 (2014)

53. Vaupel, P., Höckel, M., Mayer, A.: Detection and characterization of tumor hypoxia using pO2 histography. Antioxid Redox Signal 9(8), 1221–1235 (2007)

54. Talks, K.L., Turley, H., Gatter, K.C., Maxwell, P.H., Pugh, C.W., Ratcliffe, P.J., Harris, A.L.: The expression and distribution of the hypoxia-inducible factors HIF-1alpha and HIF-2alpha in normal human tissues, cancers, and tumor-associated macrophages. Am J Pathol 157(2), 411–421 (2000)

55. Safran, M., Kim, W.Y., O’Connell, F., Flippin, L., Günzler, V., Horner, J.W., Depinho, R.A., Kaelin, W.G.: Mouse model for noninvasive imaging of HIF prolyl hydroxylase activity: assessment of an oral agent that stimulates erythropoietin production. Proc Natl Acad Sci U S A 103(1), 105–110 (2006)

56. Bhandari, V., Hoey, C., Liu, L.Y., Lalonde, E., Ray, J., Livingstone, J., Lesurf, R., Shiah, Y.J., Vujcic, T., Huang, X., Espiritu, S.M.G., Heisler, L.E., Yousif, F., Huang, V., Yamaguchi, T.N., Yao, C.Q., Sabelnykova, V.Y., Fraser, M., Chua, M.L.K., van der Kwast, T., Liu, S.K., Boutros, P.C., Bristow, R.G.: Molecular landmarks of tumor hypoxia across cancer types. Nature Genetics 51(2), 308–318 (2019). doi:10.1038/s41588-018-0318-2

57. Schödel, J., Klanke, B., Weidemann, A., Buchholz, B., Bernhardt, W., Bertog, M., Amann, K., Korbmacher, C., Wiesener, M., Warnecke, C., Kurtz, A., Eckardt, K.U., Willam, C.: HIF-prolyl hydroxylases in the rat kidney: physiologic expression patterns and regulation in acute kidney injury. Am J Pathol 174(5), 1663–1674 (2009)

58. Rosenberger, C., Mandriota, S., Jürgensen, J.S., Wiesener, M.S., Hörstrup, J.H., Frei, U., Ratcliffe, P.J., Maxwell, P.H., Bachmann, S., Eckardt, K.U.: Expression of hypoxia-inducible factor-1alpha and -2alpha in hypoxic and ischemic rat kidneys. J Am Soc Nephrol 13(7), 1721–1732 (2002)

