## Supplementary file legends for "Hypoxia classifier for transcriptome datasets"

**Figure S1 Identification and characterization of faulty classification trees in clear cell renal carcinoma samples. A**: Representative example of a poorly performing tree. **B-D**: Ranking percentiles of genes common in poorly performing trees in the samples making up the clear cell renal carcinoma validation set. The red line represents the mean split point for each gene in the poorly performing trees. **B:** ANKDR37. **C:** NDRG1. **D:** BHLHE40.

**Figure S2 Unsupervised clustering matches areas classified as hypoxic**. UMAP representations (first column) and microscopy overlay (second column) of spot clustering using a shared nearest neighbor (SNN) based algorithm. Spots marked as hypoxic by the decision trees group together and "hypoxic" clusters are positioned close by in the UMAP space. **A**: Human Prostate Cancer, Adenocarcinoma with Invasive Carcinoma (FFPE). **B**: Adult Mouse Kidney (FFPE). **C**: Human Glioblastoma. **D**: Human Colorectal Cancer.

**Figure S3 Hypoxic genes DE between Visium dataset clusters**. -$\text{Log}_{10}$( FDR adjusted p-values ) of genes significantly up-regulated between clusters, according to their up-regulation status in the previously cited hypoxia meta-analysis. **A**: Prostate cancer, C7 vs C0. **B**: Prostate cancer, C11 vs C2. **C**: Mouse kidney, C3 vs C1. **D**: Mouse kidney, C5 vs C6.**E**: Glioblastoma, C12 vs C8.**F**: Glioblastoma, C1 vs C5 . **G**: Colorectal cancer, C14 vs C2. **H**: Colorectal cancer, C7 vs C8.

### Additional Files

Table S1

**Classification tree validation**. **Sheet 1**: metadata for each of the experiments used in the validation process. **Sheet 2**: summary of tree variables and validation accuracy. **Sheets 3-6**: performance measurements for each human validation dataset. **Sheet 7**: Proportion of TCGA tumor samples classified as hypoxic by primary site. **Sheet 8**: performance measurements for the mouse validation dataset.

Table S2

**Differential expression in unsupervised clustering of spatial gene expression datasets**: Differentially expressed genes between clusters containing a high proportion of spots classified as hypoxic and clusters with low or no hypoxic spots. **Sheet 1**: Prostate cancer, C7 vs C0. **Sheet 2**: Prostate cancer, C11 vs C2. **Sheet 3**: Mouse kidney, C3 vs C1. **Sheet 4**: Mouse kidney, C5 vs C6.**Sheet 5**: Glioblastoma, C12 vs C8.**Sheet 6**: Glioblastoma, C1 vs C5 . **Sheet 7**: Colorectal cancer, C14 vs C2. **Sheet 8**: Colorectal cancer, C7 vs C8. **Sheet 9**: Mann–Whitney test results for differences in p-values among genes up-regulated between clusters according to their up-regulation status in our hypoxia meta-analysis (random effect>0.7, FDR<0.01).

Table S3

**Comparison between tree classifiers and previously published hypoxia gene signatures**. Application of the method described in [1] of 8 hypoxic gene signatures to the validation datasets. Each table includes, for every sample, the hypoxic score derived from each signature, as well as the probability of being hypoxic assigned by our ensemble of the 20 trees with higher $F_1$-score. **Sheet 1:** PRJNA561635 time series. **Sheet 2:** specific RNA fractions other than total mRNA. **Sheet 3:** ccRCC tumor and healthy adjacent tissue samples. **Sheet 4:** murine datasets.

**References**

1. Bhandari, V., Hoey, C., Liu, L.Y., Lalonde, E., Ray, J., Livingstone, J., Lesurf, R., Shiah, Y.J., Vujcic, T., Huang, X., Espiritu, S.M.G., Heisler, L.E., Yousif, F., Huang, V., Yamaguchi, T.N., Yao, C.Q., Sabelnykova, V.Y., Fraser, M., Chua, M.L.K., van der Kwast, T., Liu, S.K., Boutros, P.C., Bristow, R.G.: Molecular landmarks of tumor hypoxia across cancer types. Nature Genetics **51**(2), 308–318 (2019). doi:10.1038/s41588-018-0318-2
